# Dynamic early recruitment of GAK-Hsc70 regulates coated pit maturation

**DOI:** 10.1101/2025.02.18.638964

**Authors:** Zhangping He, Peiyao Zuo, Peiliu Xu, Haozhi Yuan, Madhura Bhave, Xiangying Wei, Ziyan Yang, Lu Han, Sandra L. Schmid, Zhiming Chen

## Abstract

Clathrin-mediated endocytosis (CME) is the process by which clathrin assembles on the plasma membrane to form clathrin-coated pits (CCPs), which then invaginate, accumulate cargo and are released by fission from the membrane to form clathrin-coated vesicles (CCVs). A transition of nascent CCPs from flat-to-curved has been observed by various methods. However, what drives this transition remains unknown and controversial. GAK and its chaperone protein, Hsc70, are well-known to mediate clathrin release from CCVs and several studies have observed a late burst of GAK recruitment as CCVs form. Other studies have proposed that early recruitment of GAK-Hsc70 could function to provide the necessary energy source to remodel nascent flat clathrin lattices, replacing hexagons with pentagons and enabling a gain of curvature and invagination of the growing CCP; however, direct functional evidence is lacking. Here we show that GAK knockdown inhibits CCP formation and invagination. Furthermore, mutations in the J domain of GAK that abolish Hsc70 recruitment to and activation at CCPs, lead to the accumulation of GAK at CCPs, hinder CCP stabilization and invagination and result in a striking increase in the proportion of short-lived, abortive CCPs. These findings support the hypothesis that GAK-Hsc70 promotes the turnover and remodeling of nascent clathrin assemblies required for curvature development during CME.

**Significance Statement:** GAK and its chaperone protein, Hsc70, are known to be recruited to clathrin-coated vesicles (CCVs) to mediate clathrin uncoating. Previous studies have proposed that early recruitment of GAK-Hsc70 to CCPs could function to remodel nascent flat clathrin lattices, replacing hexagons with pentagons and enabling a gain of curvature of the assembling polymeric coat. However, there are conflicting views, and direct functional evidence is lacking. Here we show that GAK knockdown inhibits CCP formation and invagination. A detailed domain-specific mutational analysis of GAK pinpointed the importance of J domain-Hsc70 interactions in regulating crucial early steps of CME including CCP stabilization and invagination. These findings support a hypothesis that GAK-Hsc70 promotes turnover of clathrin at nascent CCPs required for curvature development.

## Introduction

Clathrin-mediated endocytosis (CME) is an essential cellular process that facilitates the internalization of diverse cargo molecules, including nutrients, signaling receptors, transmembrane ion channels and transporters (1–4). Dysregulation of CME has been implicated in various diseases, including cancer, neurodegenerative and cardiovascular disorders (5–8). CME is a highly orchestrated pathway initiated by the assembly of clathrin triskelions into a lattice structure at the plasma membrane (PM) leading to formation of the clathrin-coated pits (CCPs). These pits undergo a series of dynamic morphological changes, including invagination, closure, and eventual scission from the membrane, resulting in the formation of clathrin-coated vesicles (CCVs) (1, 4, 9). Live cell Total Internal Reflection Fluorescence Microscopy (TIR-FM) studies have revealed that a significant subset of nascent CCPs, termed abortive CCPs, fail at different stages of the maturation process and rapidly disassemble. A critical stage required for the formation of productive pits is early curvature generation and invagination of the growing CCP (10–12). Short-lived abortive pits are typically dim and characteristically flat (10, 12, 13).

Electron microscopy studies have revealed that flat clathrin lattices are composed almost entirely of hexagonal arrays, while curved lattices consist of both hexagons and pentagons (14–18). Some intermediate structures have been observed and proposed to represent the transition from hexagons to pentagons needed to generate curvature (11, 14), although this has been a matter of debate (19–23). While the structural transitions such as the flat-to-curved transformation of the clathrin lattice have been observed using various experimental approaches (20–25), the molecular mechanisms driving these transitions remain poorly understood.

The ATP-dependent chaperone Hsc70 (Heat shock cognate protein 70) and its co-chaperones auxilin and GAK (cyclin G-associated kinase, also known as auxilin 2) are well-known for their roles in the uncoating of CCVs, a critical late step in CME that allows clathrin and associated adaptor proteins to be recycled for subsequent rounds of endocytosis (4, 26, 27). Mechanistically, auxilin and GAK recruit Hsc70 to the clathrin lattice and stimulate its ATPase activity providing the energy required to disrupt the interactions between clathrin triskelions and disassemble the clathrin coat (4, 28–32). Consistent with its role in the uncoating reaction, several studies have reported a burst of GAK recruitment to CCPs at late stages, concurrent with CCV formation and uncoating (33–35).

Other studies, however, have reported roles for GAK during early stages of CME (34, 36–38). and have observed smaller, fluctuating bursts of GAK appearance throughout the lifetimes of CCPs (33, 36). One proposed role for the early recruitment of GAK-Hsc70 is to provide the energy necessary to remodel the clathrin lattice, replacing hexagonal arrangements with pentagons to enable curvature formation (4, 11, 14). Despite this intriguing hypothesis, direct functional evidence supporting such a role for GAK-Hsc70 has been lacking.

Auxilin and GAK are multidomain proteins that both encode a clathrin and AP2-binding domain, a J domain that recruits and activates Hsc70, and a PTEN-like domain that binds phosphatidylinositol lipids. GAK also encodes an N-terminal kinase domain that phosphorylates the µ2 subunit of AP2 (38). Interestingly, in two studies assessing the effects of knockdown of endocytic accessory factors on CCP dynamics (39, 40), GAK has emerged as an outlier exhibiting multiple, diverse phenotypes (36).

In this study, we confirm and further characterize a critical early role for GAK in CME. By selectively mutating each of GAK’s functional domains, we dissect their roles in CME and CCP dynamics. Surprisingly, only mutations in the clathrin-binding and J domain have strong effects on CME and CCP dynamics. We demonstrate that GAK is essential for CCP maturation and that its ability to recruit Hsc70 is critical for this process. Disruption of the GAK-Hsc70 interaction results in the accumulation of GAK at CCPs, impaired CCP initiation, stabilization and invagination, and rapid turnover of clathrin assemblies. These findings provide new insights into the molecular mechanisms underlying clathrin lattice remodeling and support the hypothesis that the GAK-Hsc70 complex promotes the turnover of clathrin triskelions required for flat-to-curved transition of clathrin assembly during CME.

## Results

### GAK knockdown inhibits TfnR uptake and CCP formation

To explore the functions of GAK in CME, we first examined the effects of siRNA-mediated knockdown of GAK on CME cargo uptake and CCP dynamics in ARPE19-HPV16 cells that stably express eGFP-CLCa (herein called ARPE-HPV eGFP-CLCa) (41). The expression of GAK was efficiently knocked down by transfection of siRNA directed towards its 3’UTR (Fig. 1A), resulting in significant accumulation of TfnR, a prototypic cargo of CME (42), on cell surface (Fig. 1B, Fig. S1), and strong reduction of the cellular uptake efficiency of TfnR (Fig. 1C).

**Figure 1.**
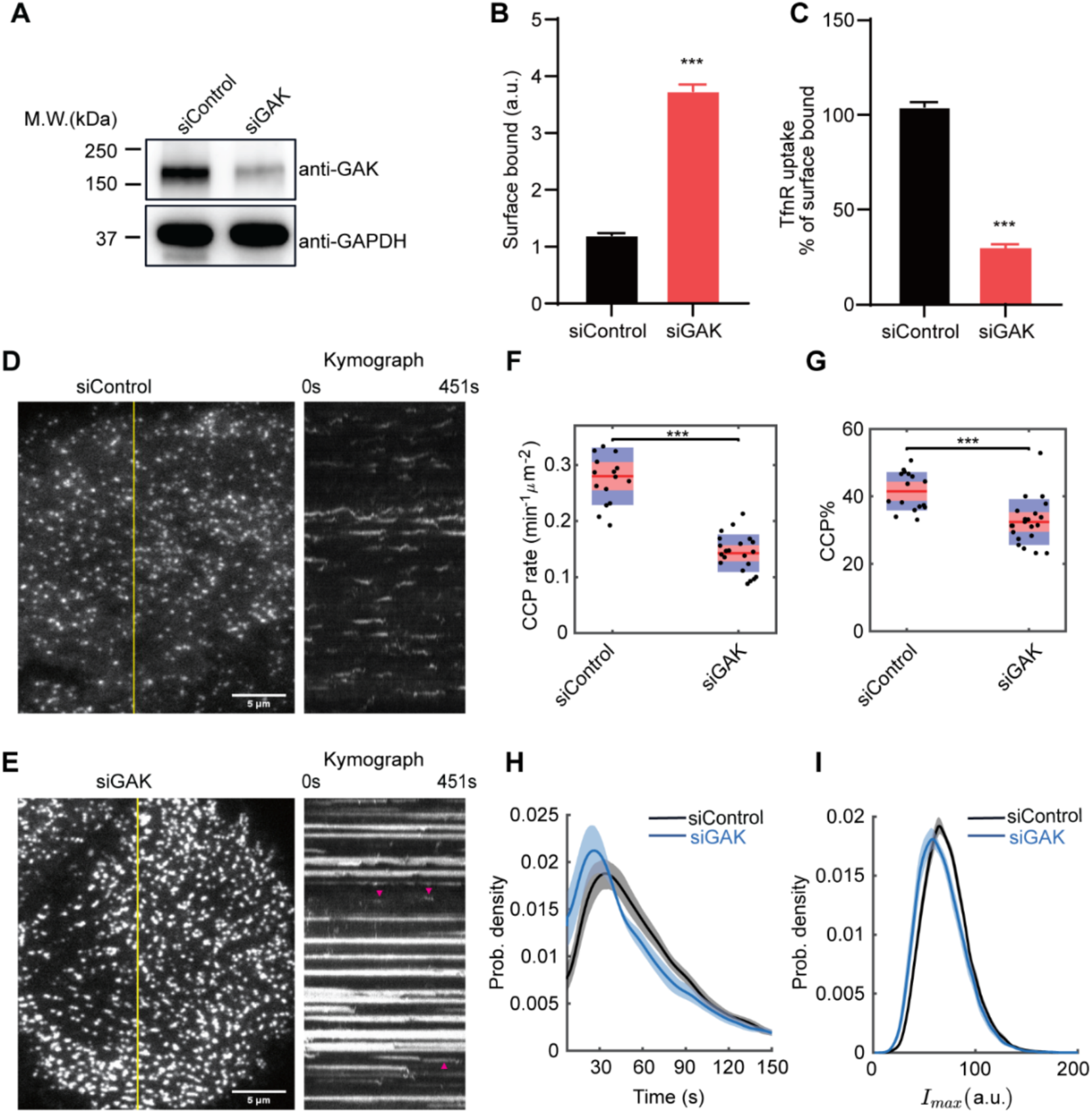
GAK knockdown inhibits TfnR uptake and *bona fide* CCP formation. **(A)** siRNA-mediated knockdown of GAK in ARPE-HPV eGFP-CLCa cells. **(B-C)** Surface bound and uptake efficiency (Internalized/Surface-bound) of TfnR measured by in-cell ELISA at 10min. Error bars: SEM of n = 8 samples. Statistical analysis of the data in (B-C) was performed using GraphPad Prism 8 by unpaired t-test: ***, P ≤ 0.001. **(D-E)** Representative single frame images from TIRFM movies (7.5 min/movie, 1 frame/s, see Movie 1) and corresponding kymographs from region indicated by yellow lines of ARPE-HPV eGFP-CLCa cells treated with (D) control siRNA or (E) GAK siRNA. Red arrowheads point to dynamic structures. Scale bars = 5µm. **(F-G)** Effect of GAK knockdown on (F) the initiation rate and (G) the % of bona fide CCPs. Each dot represents a movie. Statistical analysis of the data in (F) and (G) is the Wilcoxon rank sum test, ***, P ≤ 0.001. **(H-I)** Distributions of (H) the lifetime and (I) the maximum fluorescence intensity of bona fide CCPs. Data presented were obtained from a single experiment (N = 15 movies for each condition) that is representative of 3 independent repeats. Number of dynamic tracks analyzed: 184,845 for siControl and 138,729 for siGAK. Shadowed area in (H,I) indicates 95% confidential interval.

To further define which stages of CME were dependent on GAK, we used quantitative live-cell TIR-FM to visualize clathrin-coated structures (CCSs) and analyze CCP dynamics (43) in GAK-depleted ARPE-HPV eGFP-CLCa cells. GAK knockdown resulted in the formation of brighter and static CCSs (Fig. 1D,E & Movie 1), which have been observed under other perturbation conditions (12, 41) and reflect the accumulation of CCPs that are either larger, flatter or both. However, we also noted a subpopulation of dynamic CCPs (red arrowheads, Fig. 2E) that were visually obscured by the bright static CCSs. Indeed, despite the deceptive nature of the images, the percentage of static CCSs (lifetime > 150s) was only 18%, as determined by unbiased quantitative analysis.

**Figure 2.**
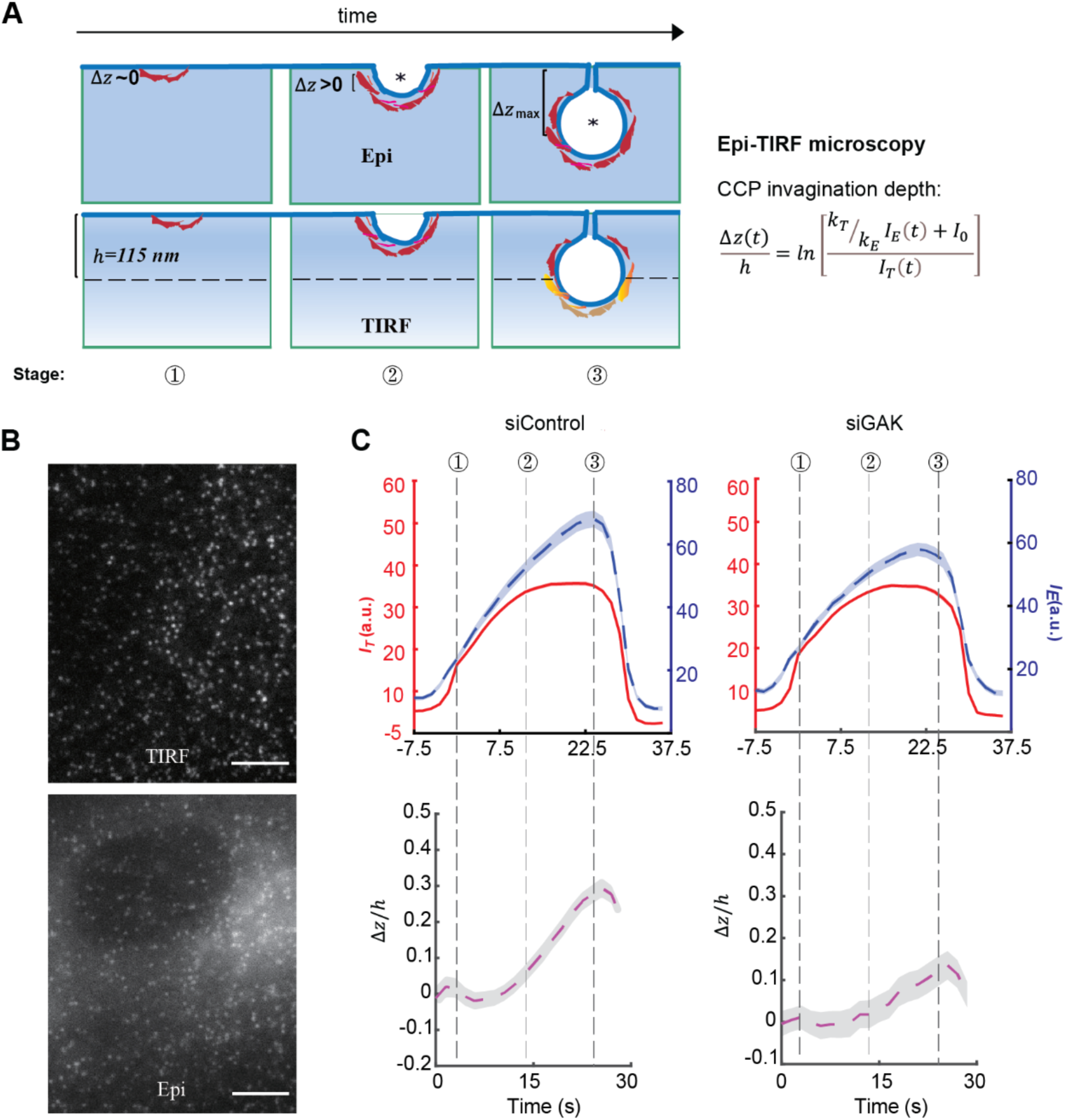
GAK knockdown inhibits CCP invagination. **(A)** Scheme of Epi-TIRF microscopy for measuring the invagination of CCPs using primary/subordinate tracking. Δ*z*(*t*) denotes the invagination depth of CCPs over time, while *I_E_* and *I_T_*denotes the cohort-averaged fluorescence intensity from Epi and TIRF channels, respectively. *k_E_* and *k_T_* are the initial growth rate for the Epi and TIRF channel signals, respectively. *I*_0_ is an additive correction factor. ‘*’ indicates the mass center of clathrin coat. *h* = 115nm is the evanescent depth of TIRF field, see more detail in ref. (10). **(B)** Representative TIRF and Epi microscopy images. Scale bars = 10 µm. **(C)** Top: Cohort-averaged CCP fluorescence intensity traces from Epi (blue) and TIRF (red) channels; bottom: calculated Δ*z*(*t*) curves. Data presented were obtained from n = 16 movies for each condition. Number of CCP tracks analyzed to obtain the Δ*z*/*h* curves: 14355 for siControl and 5330 for siGAK. Shadowed area in (C) indicates 95% confidential interval.

To quantify the dynamic behaviors of CCPs we performed cmeAnalysis (12, 44, 45) and DASC (disassembly asymmetry score classification) (10), which together provide a comprehensive and unbiased characterization of CCP intermediates and CME progression. These analyses revealed that GAK knockdown significantly reduced the initiation rate (Fig. 1F) and percentage of *bona fide* CCP (Fig. 1G), thus resulting in a corresponding decrease in the lifetimes (Fig 1H) and intensity (Fig. 1l) of CCPs, both indicators of an increase in abortive CCPs (10).

Together these results are consistent with previous reports of an early role for GAK in CME (39, 40), and provide evidence that GAK regulates CCP initiation, stabilization and maturation.

### GAK knockdown inhibits CCP invagination

Successful invagination of the clathrin coat has been identified as a key process that determines the fate of CCPs as either productive or abortive (10). To determine whether GAK regulates CCP invagination, we conducted Epifluorescence (Epi)-TIRF microscopy to measure CCP invagination in live cells (Fig. 2A). In this approach, time-lapse Epi and TIR fluorescence signals were near-simultaneously acquired for ARPE-HPV eGFP-CLCa cells treated with control or GAK siRNA (Fig. 2B). Acquired imaging data was analyzed using cmeAnalysis with TIRF as a primary channel and Epi as a secondary channel. The resulting Epi and TIR fluorescence intensity traces of CCPs with similar lifetimes were then aligned and averaged by DASC to yield intensity cohorts that were further log-transformed to give average traces of the invagination depth (Δ*z*) of the CCPs’ center-of-mass (Fig. 2C). For this analysis we chose to present the ~30s lifetime cohort because they represent the invagination behavior of the most frequent tracks (Fig. 1H). In control cells, curvature acquisition was delayed relative to clathrin assembly, as has been previously observed (10, 13). GAK knockdown strongly inhibited CCP invagination (Fig. 2C). Importantly, this effect and all other phenotypes associated with GAK knockdown were fully rescued by expression of siRNA resistant GAK-mRuby(WT) (Fig. S2 & Movie 1). Interestingly, siRNA-mediated knockdown of auxilin did not affect CCP initiation, stabilization or invagination in ARPE-HPV cells (Fig. S3).

Together these data suggest a key early role for the ubiquitously expressed GAK, but not the neuron-enriched auxilin, in CME in regulating curvature generation at nascent CCPs.

### Hsc70 recruitment mediated by the J domain of GAK is essential for CME regulation

Next, we explored the mechanism by which GAK regulates CME. GAK is a multi-domain protein that consists of an N-terminal kinase domain (35, 46), a PTEN-like domain (34), a clathrin/AP2-binding domain (47), as well as a C-terminal J domain (46) (Fig. S4A). The N-terminal kinase domain of GAK can phosphorylate clathrin accessory proteins, such as the μ2 subunit of AP1 and AP2 (38). The PTEN-like domain of GAK binds preferentially to phosphatidylinositol monophosphates (37) and may act as a switch for uncoating clathrin-coated vesicles (CCVs) (34, 35). The clathrin/AP2 domain contains binding sites for clathrin and AP2 (47). The J domain contains an HPD motif that recruits and activates Hsc70 to facilitate the uncoating of clathrin-coated vesicles (38, 48). To explore which domain function(s) might contribute to the regulation of CME by GAK, we generated a series of GAK constructs containing loss-of-function mutations in each domain (denoted as Kinase*, PTEN*, clathrin*, AP2* and J*, respectively) (37, 38, 49–51), and then stably expressed the corresponding mutants with C-terminal tagged eGFP or mRuby in ARPE-HPV mRuby-CLCa or ARPE-HPV eGFP-CLCa cells, respectively (Fig. S4). All fluorescently tagged cell lines were FACS-sorted to achieve homogenous expression levels that were ~2-fold above endogenous levels.

To examine the effects of domain-specific loss-of-function mutations in CME, 3’ UTR siRNA was used to knockdown endogenous but not exogenous GAK (Fig. S4B,C). We then assessed TfnR uptake (52, 53), as well as CCP dynamics using live-cell TIR-FM and Epi-TIRF microscopy to quantitatively evaluate changes in CME. Although previous studies had shown that transient overexpression of WT GAK inhibited Tfn uptake by sequestering clathrin (54), the much lower levels of GAK expression in our stably transfected cell lines had no effect (Fig. 3A,B, Fig. S2). Interestingly, loss-of-function mutations in the kinase domain (Kinase*), PTEN-like domain (PTEN*), and mutations that abolish AP2-binding ability (AP2*) did not affect TfnR uptake (Fig. 3A,B) or the dynamic behavior of CCPs (Movie 2), including the initiation rate (Fig. 3C), proportion (CCP%, Fig. 3D), lifetime distribution (Fig. 3E), maximum fluorescence intensity distribution (Fig. 3F), or the invagination depth (Fig. 3G) of *bona fide* CCPs. These results suggest that the respective activities of kinase domain, PTEN-like domain, and AP2-binding motifs are not required for the regulation of CME by GAK.

**Figure 3.**
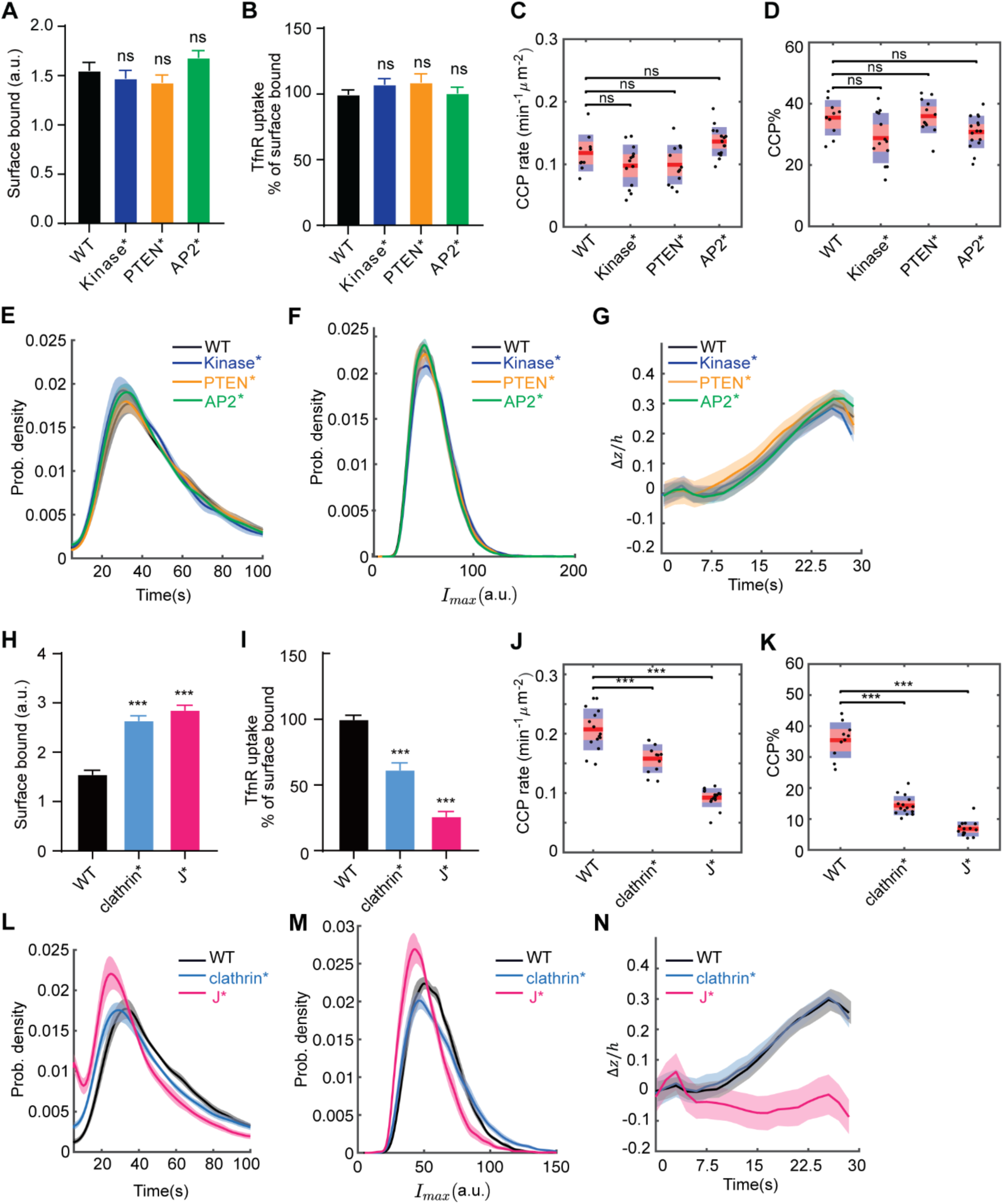
A GAK J domain mutant that abolishes GAK-Hsc70 interactions inhibits CCP formation and invagination. **(A-G)** Effects of Kinase*, PTEN*, and AP2* on (A-B) TfnR uptake, (C-F) CCP formation and (G) CCP invagination. Number of dynamic tracks analyzed in (C-F): 91,648 for eGFP-GAK(WT), 91,535 for eGFP-GAK(Kinase*), 121,967 for eGFP-GAK(PTEN*) and 159,505 for eGFP-GAK(AP2*). Number of CCP tracks analyzed to obtain the Δ*z*/*h* curves in (G): 11,204 for WT, 13,934 for Kinase*, 8,501 for PTEN*, 15,464 for AP2*. **(H-N)** Effects of CBD* and J* mutants on (H-I) TfnR uptake, (J-M) CCP formation and (N) CCP invagination. Number of dynamic tracks analyzed in (J-M): 91,648 for eGFP-GAK(WT), 156,202 for eGFP-GAK(CBD*), and 509,429 for eGFP-GAK(J*). Number of CCP tracks analyzed to obtain the Δ*z*/*h* curves in (N): 11,204 for WT, 7,198 for CBD*, 3,888 for J*. Cell lines used in (A-F) and (H-M): ARPE-HPV mRuby-CLCa+GAK(WT or mutant)-eGFP. Cell lines used in (G) and (N): ARPE-HPV eGFP-CLCa+GAK-mRuby. Error bars in (A,B,H,I): SEM of n = 8 samples. Statistical analysis of the data in (A,B,H,I) was performed using GraphPad Prism 8 by unpaired t-test: ns, not significant; ***, P ≤ 0.001. Each dot in (C,D,J,K) represents a movie. Statistical analysis of the data in (C,D,J,K) is the Wilcoxon rank sum test: ns: not significant, P > 0.05; ***, P ≤ 0.001. Shadowed area in (E-G) and (L-N) indicates 95% confidential interval.

Importantly, mutations that impair clathrin-binding (clathrin*, Fig. S5) and Hsc70 recruitment (J*) significantly increased surface-bound TfnR (Fig. 3H) and correspondingly reduced TfnR uptake efficiency (Fig. 3I). In addition, both clathrin* and J* mutants altered the dynamic behaviors of CCSs (Movie 3). Significant reduction in the initiation rate (Fig. 3J) and proportion (CCP%, Fig. 3K) of *bona fide* CCPs were observed for clathrin* and J*, indicating that both clathrin-binding and Hsc70 recruitment are important for GAK to regulate CCP initiation and stabilization. Moreover, J*, but not clathrin*, dramatically shortened the lifetime (Fig. 3L & Movie 3), strongly reduced the maximum fluorescence intensity (Fig. 3M), and substantially inhibited the invagination (Fig. 3N) of CCPs. Together, these results reveal a striking role for GAK’s J domain-mediated recruitment of Hsc70 in regulating CCP initiation, stabilization and invagination.

### Inhibiting GAK-Hsc70 interactions with MAL3-101 phenocopies the expression of GAK(J*)

To confirm that Hsc70 recruitment by GAK is crucial in CME regulation, we tested the effect of MAL3-101, a small molecule inhibitor of the interactions between the J domain of GAK and Hsc70 (55) on CME. ARPE-HPV eGFP-CLCa cells were treated with 10 µM MAL3-101 for 3h during which no obvious effects on cell viability, proliferation, or cytotoxicity were observed (Fig. 4A). Treatment with MAL3-101 significantly reduced the initiation rate (Fig. 4B), proportion (CCP%, Fig. 4C), lifetime (Fig. 4D), maximum fluorescence intensity (Fig. 4E), and invagination depth (Fig. 4F) of *bona fide* CCPs, exactly mirroring the effects of the J* mutation. Together these data demonstrate that recruitment and activation of Hsc70 by the J domain is crucial for the early-stage regulation of CME by GAK.

**Figure 4.**
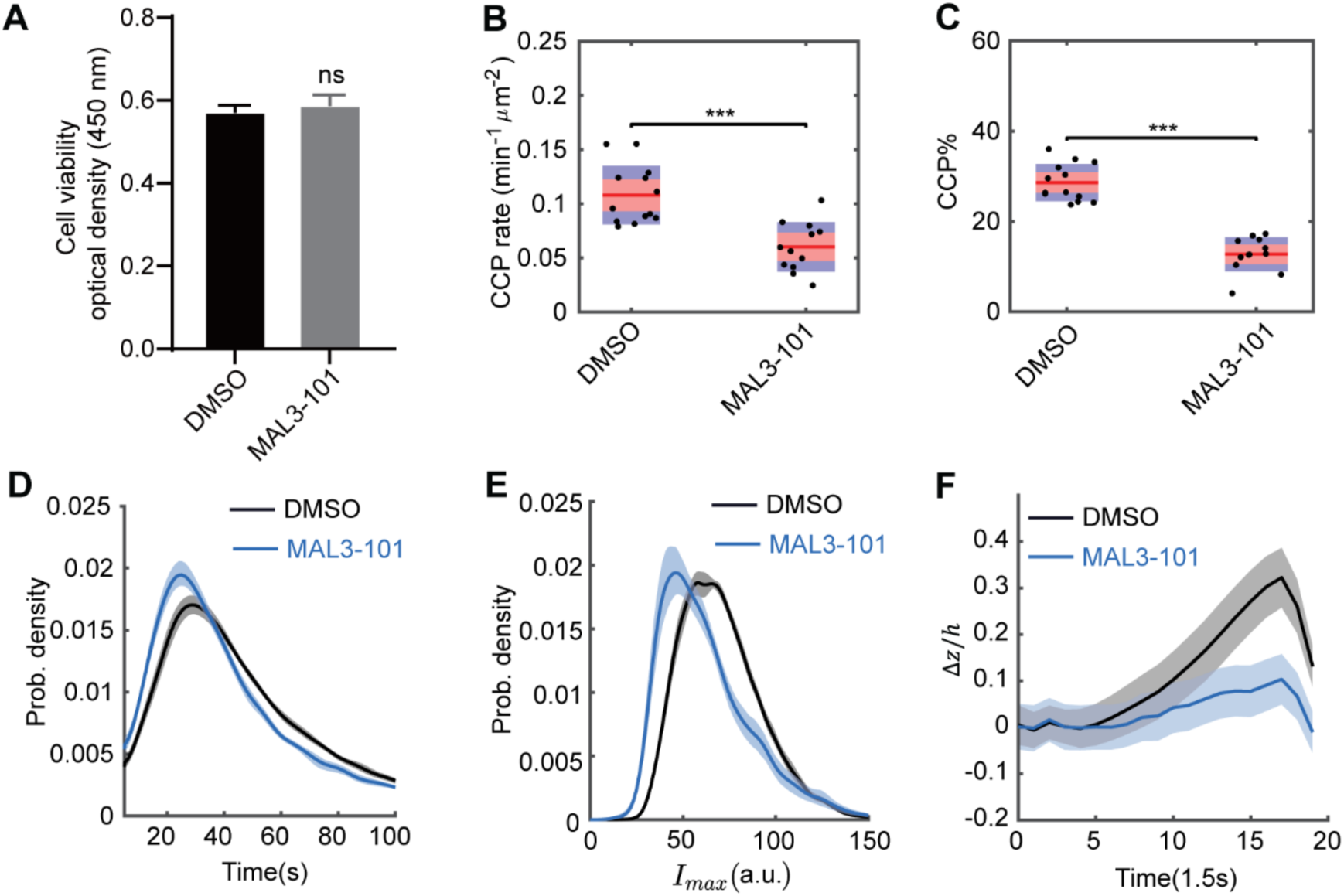
Inhibiting GAK-Hsc70 interactions with MAL3-101 phenocopies the effects of GAK(J*). **(A) Cell viability measured by** CCK8 assay of ARPE-HPV eGFP-CLCa cells treated with 10 uM DMSO or MAL3-101 for 3h. Error bars: SEM of N = 8 samples. Statistical analysis of the data in (A) was performed using GraphPad Prism 8 by unpaired t-test: ns, not significant, P > 0.05. **(B-E)** ARPE-HPV eGFP-CLCa cells were treated with 10 µM MAL3-101 for 3h, resulting in reduced (B) initiation rate, (C) %, (D) lifetime and (E) maximum fluorescence intensity of bona fide CCPs. Number of dynamic tracks analyzed: 76,963 for DMSO, 131,301 for MAL3-101. Statistical analysis of the data in (B,C) is the Wilcoxon rank sum test: ***, P ≤ 0.001. **(F)** Treatment of MAL3-101 strongly inhibited CCP invagination. Number of CCP tracks analyzed to obtain the Δ*z*/*h* curves in (F): 11,204 for DMSO and 8,051 for MAL3-101. Shadowed area in (D-F) indicates 95% confidential interval.

### WT GAK is dynamically recruited to CCPs while J* accumulates with clathrin in CCPs

To gain further insight into the regulation mechanism of GAK-Hsc70 in CME, we conducted dual-channel (mRuby-CLCa and GAK-eGFP) TIR-FM imaging to directly probe the recruitment dynamics of GAK (WT, clathrin* or J*) during CME. Averaging across a representative 30s cohort of CCPs showed low levels of GAK recruitment at early stages, followed by a burst of GAK recruitment that peaked as clathrin intensity precipitously declined (Fig. 5A). A similar pattern was observed for the clathrin* mutant, although the extent of recruitment was significantly diminished (Fig. S5). Importantly, when individual tracks were analyzed, and consistent with a previous report (34), small and variable amounts of WT GAK were observed to be recruited transiently during clathrin-coat assembly and prior to the final burst of recruitment (Fig. 5B,C and see other examples in Fig. S6A). A very different pattern was observed for the J* mutant, which was co-recruited to CCPs and accumulated along with clathrin throughout their shortened (Fig. 3L) lifetimes (Fig. 5D-F & S6B). These results suggest that GAK-dependent recruitment and activation of Hsc70 are required for the fluctuating interactions between GAK and clathrin at CCPs, and in turn, that fluctuations in the recruitment of GAK and active Hsc70 during early stages of CME are required for CCP invagination and stabilization (Fig. 6).

**Figure 5.**
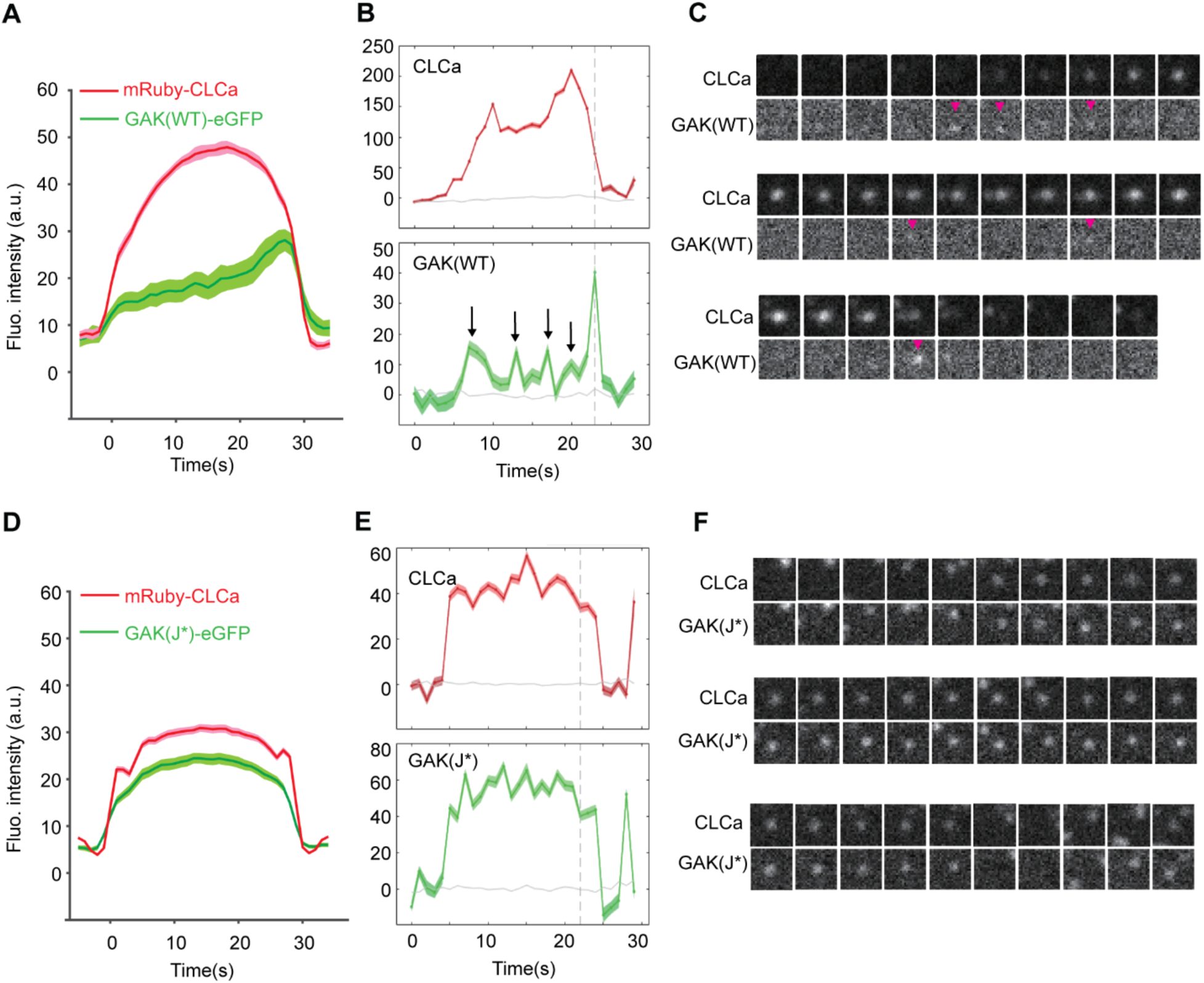
GAK(WT) is transiently recruited to growing CCPs while GAK(J*) accumulates along with clathrin at CCPs. **(A-C)** Averaged fluorescence intensity cohorts (A) and representative single track of mRuby-CLCa and GAK(WT)-eGFP (B-C). **(D-F)** Fluorescence intensity cohorts (D) and representative single track of mRuby-CLCa and GAK(J*)-eGFP (E-F).

**Figure 6.**
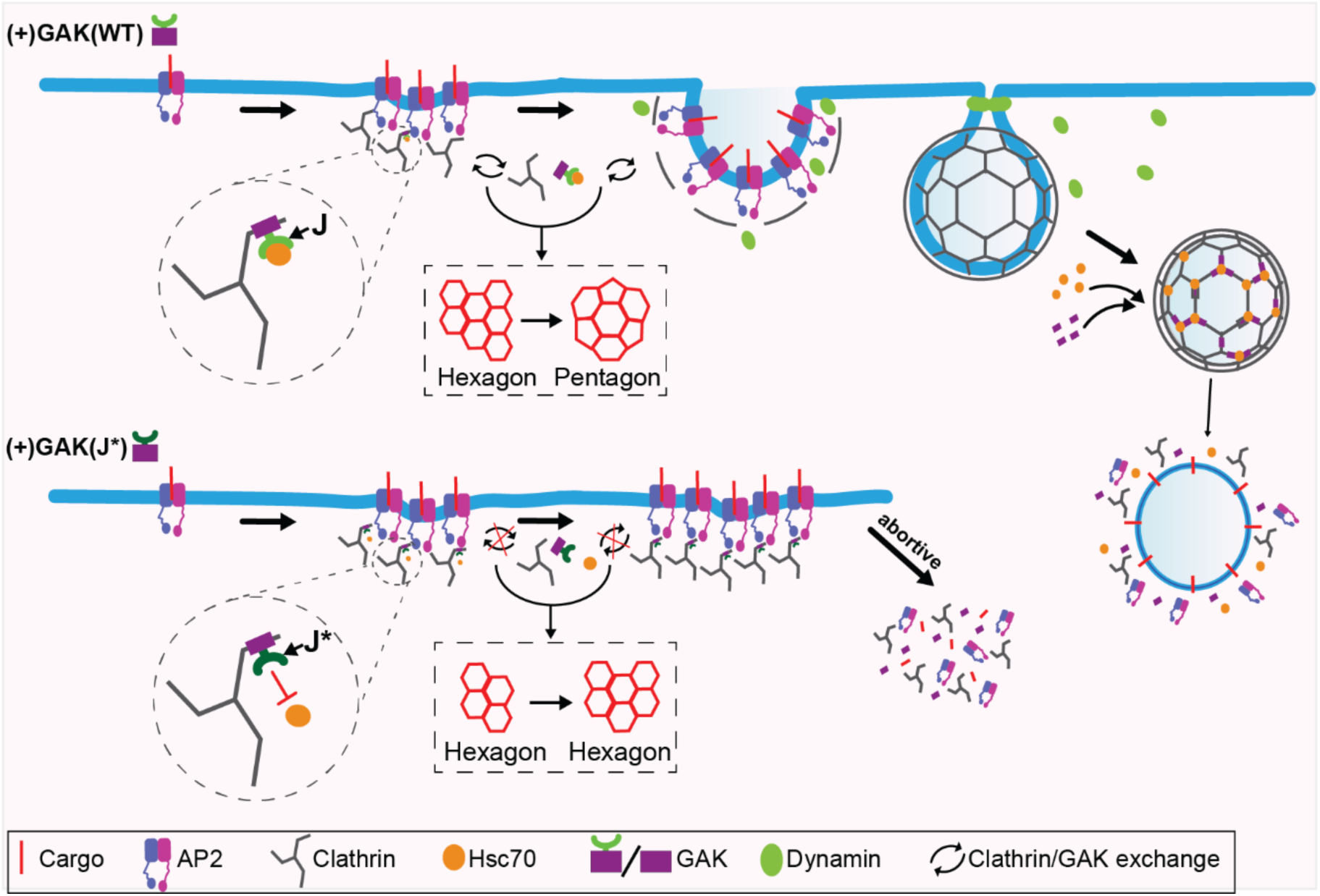
A model for GAK-Hsc70 regulation of CCP assembly and invagination. Dynamic recruitment of Hsc70 via the J domain of GAK regulates the exchange of clathrin triskelia that enables remodeling of the clathrin lattice and the development of curvature at nascent clathrin-coated pits. Abolishing early GAK-dependent Hsc70 recruitment and activation inhibits clathrin exchange and results in the accumulation of GAK, while CCPs fail to develop curvature and are quickly aborted.

## Discussion

In this study, we have uncovered a novel role for the dynamic recruitment of GAK-Hsc70 in regulating the early stages of clathrin-mediated endocytosis (CME), beyond its well-established role in clathrin uncoating (4, 26, 27). Inhibiting, either by mutagenesis or chemically, the ability of GAK’s J domain to recruit and activate Hsc70 at growing CCPs severely impairs CCP invagination and stabilization, resulting in an increase in abortive CCPs. While previous studies have reported the dynamic, ATP-dependent exchange of clathrin at nascent CCPs (15, 22), to our knowledge, our findings are the first to demonstrate a requirement for functional GAK-Hsc70 interactions to catalyze the remodeling of clathrin lattices needed to enable early curvature generation at nascent CCPs. Our results position the GAK-Hsc70 complex as a central regulator of the morphological transition from flat to curved clathrin lattices, a process essential for productive CCP formation.

### Mechanistic insights into GAK’s function in CME

GAK is a multidomain protein and our functional dissection of GAK domains, coupled to real-time imaging, provided further insights into GAK activity in CME and regulating CCP dynamics. We showed that GAK’s clathrin-binding domain plays a critical role in enabling the dynamic remodeling of clathrin lattices during CCP initiation and stabilization. Loss-of-function mutations that abolish GAK’s ability to bind clathrin led to severe defects in CCP formation and maturation and an increase in abortive CCPs. In contrast to the clathrin-binding and J-domains, we were unable to detect any effects on CME or CCP dynamics by inactivating mutations of the kinase domain, PTEN-like domain, or AP2-binding motifs. We speculate that loss of GAK kinase activity could be compensated for by redundant activities of other µ2 kinases such as AAK1 (56), while loss of AP2 binding might reflect a greater role for clathrin binding in recruiting GAK to CCPs. Finally, loss of lipid binding by the PTEN-like domain is consistent with the suggestion that this activity plays a later role in triggering rapid CCV uncoating (35). Importantly, these observations are in line with a previous report (46) in which expression of a fragment of GAK containing only the clathrin-binding and J-domains could fully rescue the CME and behavior defects caused by GAK knockout in fibroblasts and in mice.

### GAK-Hsc70 acts as a dynamic remodeler of the clathrin lattice

Consistent with our findings, previous live cell imaging studies of GAK recruitment to CCPs have observed a burst of GAK recruitment at late stages, coincident with the acute disappearance of clathrin and presumably indicative of the onset of CCV uncoating (33, 34, 37). However, results regarding earlier and transient recruitment of GAK to growing CCPs are conflicting. He et al., (35) could only detect the late burst of recruitment of genome-edited GAK-eGFP to CCPs and suggested that observations of earlier recruitment might have been an artifact of overexpression. Indeed, transient overexpression of GAK inhibits CME by sequestering clathrin (54). Here we have selected stable cell lines expressing modest levels of GAK-eGFP that have no effect on CME. Moreover, we provide functional evidence of an early role for GAK in CCP stabilization that is dependent on its ability to bind clathrin and recruit and activate Hsc70. Finally, we provide evidence that Hsc70 activation drives the release of GAK from CCPs, accounting for its transient association during CCP growth and maturation. Our observation that the J domain mutant of GAK is recruited to and accumulates at CCPs coincidently with clathrin growth is consistent with its ability to be recruited to and function at nascent CCPs.

Interestingly, while mutating the clathrin binding domains of GAK severely inhibit CCP stabilization, it has no effect on CCP invagination. Our data suggest that in the absence of clathrin binding motifs, the remaining AP2 binding motifs are sufficient to recruit small amounts of GAK and Hsc70 sufficient to remodel clathrin lattices for curvature generation. Indeed, biochemical studies have shown that very small amounts of GAK are needed to catalyze clathrin release (57). That the clathrin-binding mutant nonetheless severely inhibits CCP stabilization suggests a structural role for GAK’s clathrin binding domain in stabilizing the nascent clathrin lattice. Indeed, GAK’s clathrin binding domain has been shown to exhibit clathrin assembly activity *in vitro* (47).

The dynamic recruitment pattern of GAK, characterized by transient early associations followed by a strong burst during uncoating, suggests precise temporal regulation of its activity. During early stages, the GAK-Hsc70 complex facilitates CCP formation and invagination by promoting dynamic exchange of clathrin triskelions, enabling the structural transitions required for membrane invagination. Later, during vesicle formation, the same molecular machinery catalyzes the complete disassembly of the clathrin coat. What prevents GAK-Hsc70 from completely disassembling nascent CCPs? We propose that the observed late-stage generation of phosphatidylinositol-4-phosphate, a GAK PTEN-like domain substrate (58) stabilizes GAK at CCPs, even in the midst of Hsc70 uncoating activity. Indeed, the PTEN-like domain is required for the late burst of GAK recruitment to CCPs (34). This differential regulation of early vs late recruitment of GAK ensures that early GAK-Hsc70 activity promotes lattice remodeling without causing premature coat disassembly, while the later burst drives complete uncoating. This dual functionality represents an elegant example of how cells can utilize a single protein to achieve different functionality by repurposing the same molecular machine for distinct but mechanistically related tasks.

Interestingly, nascent CCPs that fail to gain curvature in J* expressing cells are still rapidly turned over, even when GAK-Hsc70 interactions are inhibited. Thus, the mechanisms controlling rapid disassembly for flat abortive pits remain unknown. We suggest 3 possibilities: i) disassembly is mediated by auxilin-Hsc70 interactions, ii) disassembly is mediated by an as yet unknown mechanism or iii) disassembly is spontaneous. That inhibition of J domain-Hsc70 interactions with MAL3-101 phenocopies the GAK J* mutation favors possibilities ii and iii. Further studies will be needed to resolve this issue.

### A Model for GAK-Hsc70-Mediated Clathrin Lattice Remodeling

Our results suggest a model in which GAK, through its clathrin-binding domain, transiently associates with the assembling clathrin coat. The J-domain then recruits Hsc70, whose ATPase activity provides the energy required for localized clathrin disassembly and coat rearrangement. GAK is released, in an Hsc70 activity-dependent manner, together with the dissociated clathrin. This localized dynamic turnover of GAK and clathrin triskelia, avoids complete disassembly and enables the progressive remodeling of the lattice geometry, likely promoting the incorporation of pentagons into the hexagonal lattice and driving curvature generation. The subsequent dissociation of GAK-Hsc70 allows the CCP to progress towards invagination and vesicle formation (Fig. 6). At late stages, after closure of the vesicle neck and CCV formation, interactions between the PTEN-like domain of GAK with accumulating monophosphatidylinositol lipids (35, 58) stabilizes GAK allowing for complete disassembly of coat. Thus, we propose a cycle (Fig. 6) in which Hsc70 is transiently recruited to growing CCPs by GAK where it catalyzes both clathrin and GAK dissociation and then leaves bound to the dissociated clathrin. Our results thus provide direct functional evidence for a previously proposed model (11, 59) that early recruitment of GAK-Hsc70 functions to provide the necessary energy source to remodel the flat clathrin lattices, thus offering mechanistic insights into this fundamental but crucial step in CME.

## Materials and Methods

### Plasmids

A plasmid encoding GAK(WT, mouse) cDNA in a pmCherry-N1 vector backbone was obtained from Addgene (#27695). Next, site-directed mutagenesis was conducted to generate loss-of-function mutations in each domain of GAK, denoted as Kinase*, PTEN*, clathrin*, AP2* and J*, respectively. Subsequently, GAK constructs (WT and mutants) were separately cloned into a pLVx-IRES-Puro vector with a C-terminal tagged eGFP using NEBuilder® HiFi DNA Assembly Master Mix (Catalog #E2621S). Similarly, GAK constructs (WT and mutants) were also separately cloned into a pLVx-IRES-Puro-RFP670 vector with C-terminal mRuby tag. Mutagenesis and cloning primers are listed in supplementary Table S1.

### Cell culture, lentivirus infection, siRNA transfection and rescue

ARPE19-HPV16 (herein called ARPE-HPV) cells were obtained from ATCC and cultured in DMEM/F12 (Gibco, Catalog #8122502) with 10% FBS. HEK293T cells were obtained from ATCC and cultured in DMEM (Gibco, Catalog #8122070) with 10% FBS. ARPE-HPV cells that stably express eGFP-CLCa or mRuby-CLCa were generated in our previous study (41, 60).

Lentiviruses encoding GAK-eGFP were produced in HEK293T packaging cells following standard transfection protocols (61) and harvested for subsequent infections to ARPE-HPV mRuby-CLCa cells to generate ARPE-HPV mRuby-CLCa+GAK-eGFP cells. Lentiviruses encoding GAK-mRuby were produced following the same protocol and used to infect ARPE-HPV eGFP-CLCa cells to generate ARPE-HPV eGFP-CLCa+GAK-mRuby cells. All fluorescent-tagged cells were FACS sorted for homogenous mRuby and eGFP signals and passaged for 2 weeks before experiments.

For siRNA-mediated knockdown of GAK, cells were seeded on 6-well plates (250,000 cells/well) and transfected with 2 rounds of GAK 3’UTR siRNA (Mixture of hs.Ri.GAK.13.3 and hs.Ri.GAK.13.9, IDT reference #: 284315824 and 284315827) or negative control siRNA (Silencer Select Negative Control #1 siRNA, cat#:4390843) through 3 days. siRNA Transfections were mediated with Opti-MEM and Lipofectamine RNAi-MAX (Invitrogen) as detailed in ref(41). Anti-GAPDH Rabbit pAb (Abclonal, AC001 and Anti-GAK Rabbit pAb (Proteintech, 12147-1-AP were used in Western Blotting to confirm protein expression level and knockdown efficiency.

### CCK8 assay

CCK8 assay was performed using Cell Counting Kit-8 (CCK8, KeyGen BioTECH, Nanjing, China). Briefly, ARPE-HPV eGFP-CLCa cells were cultured overnight in a 96-well plate. After incubation with 10 uM DMSO or MAL3-101 for 3 hours, the supernatant was replaced with 100 µl of pre-diluted CCK8 solution (dilution ratio: 90 µl complete culture medium: 10 µl CCK8 solution) continue incubation for 2 hours. The OD values were measured at 450 nm using a microplate reader.

### Transferrin receptor (TfnR) uptake assay

In cell ELISA was used to measure TfnR uptake following established protocols described in our previous publications (41, 52, 53). Briefly, ~15,000 cells/well were seeded on gelatin-coated 96-well (1x8 stripwell) plate (Corning, #9102). After growing overnight, cells were first starved in 37°C-warm PBS4+ (1× PBS buffer plus 0.2% BSA, 1mM CaCl2, 1mM MgCl2, and 5mM D-glucose) for 30min, and then cooled down to 4°C before adding ice-cold PBS4+ containing 5µg/ml HTR-D65 (anti-TfnR mAb) (62). Next, cells were divided into three groups: 1) some cells were kept at 4°C for the measurement of surface-bound TfnR; 2) some cells were acid-washed (0.2M acetic acid and 0.2M NaCl, pH 2.3) to remove surface-bound HTR-D65 for the measurement of background signal; 3) the rest cells were first incubated in a 37°C water bath for 10min and then acid-washed (0.2M acetic acid and 0.2M NaCl, pH 2.3) for the measurement of internalized HTR-D65. All the three groups of cells were then washed with cold PBS and fixed with 4% PFA (Electron Microscopy Sciences, diluted in PBS) for 30min at 37°C. Subsequently, all cells were permeabilized with 0.1% Triton X-100 and blocked with Q-PBS (PBS, 2% BSA, 0.1% lysine, and 0.01% saponin, pH 7.4) for 2h. Surface-bound and internalized HTR-D65 were detected with HRP Goat anti-Mouse IgG (H+L) (BioRad) and o-phenylenediamine dihydrochloride (OPD, Sigma-Aldrich). Well-to-well variation of cell numbers was accounted for by BCA assays.

### TIR-FM and Epi-TIRF microscopy

Cells were seeded on gelatin-coated 35 mm glass bottom dishes (ibidi, #81218-800) overnight before live cell imaging with a Nikon Eclipse Ti2 inverted microscope equipped with: 1) an Apo TIRF/100x 1.49 Oil objective; 2) a Prime Back Illuminated sCMOS Camera (Prime BSI, 6.5 x 6.5µm pixel size and 95% peak quantum efficiency; 3) a M-TIRF module for epifluorescence (Epi) acquisition; 4) an H-TIRF module for TIRF acquisition, where penetration depth was fixed to 80nm; and 5) an Okolab Cage Incubator for maintaining 37°C and 5% CO_2_. Single-channel time-lapse TIR-FM imaging data were acquired at a frame rate of 1 frame/s. Dual-channel time-lapse TIRFM imaging data were acquired at a frame rate of 0.5 frame/s. For time-lapse Epi-TIRF imaging, Epi and TIRF images were acquired nearly simultaneously at a frame rate of 0.66 frame/s. For all time-lapse imaging, 451 consecutive images/movie were acquired, and Perfect Focus System (PFS) was applied.

The acquired data were analyzed using cmeAnalysis (12, 44) to track the lifetime and fluorescence of clathrin-coated structures. In addition, DASC (10) was applied to: 1) unbiasedly classify *bona fide* CCPs vs. abortive coats; 2) calculate CCP invagination (Δ*z*) by aligning and averaging the classified CCP tracks. Tracks that overlap with others or deviate from the properties of a diffraction-limited particle were excluded from the analysis. The software package, as well as the detailed protocol, is available at: https://github.com/DanuserLab/cmeAnalysis.

## Data availability

Original data are available from the corresponding authors upon request.

## Acknowledgement

This work is supported by the National Natural Science Foundation of China (Grant No. 32200564 to Z.C.), the Natural Science Foundation of Hunan Province, China (Grant No. 2024JJ2045 to Z.C.), and initiated with support from the National Institutes of Health grant GM73165 to S.L.S.

## Competing Interest

The authors declare no competing interest.

## Author contributions

Z.C., S.L.S. and Z.H. conceived the story and designed the experiments. Z.C., S.L.S., Z.H., M.B. and X. W. interpreted the results and wrote the manuscript with input from all authors. M.B., Z.H., and Z.Y performed the mutagenesis and molecular cloning, Z.H., P.Z., H.Y. and L.H. performed the TfnR uptake assays and the TIRF, Epi-TIRF microscopy imaging and data analysis. P. X. performed FACS sorting. P.Z. drew cartoon illustrations.

## Supplemental Information

Movie 1: Time-lapse TIRFM imaging of ARPE-HPV eGFP-CLCa cells treated with control siRNA (left) or GAK siRNA (middle), as well as ARPE-HPV eGFP-CLCa+GAK(WT)-mRuby cells that were treated with GAK siRNA (right). Images were obtained at 1 frame/s and collected for 7.5 min. Movie is accelerated 25-fold.

Movie 2: Time-lapse TIRFM imaging of ARPE-HPV eGFP-CLCa+GAK(WT)-mRuby cells (#1 from left), ARPE-HPV eGFP-CLCa+GAK(Kinase*)-mRuby cells (#2 from left), ARPE-HPV eGFP-CLCa+GAK(PTEN*)-mRuby cells (#3 from left) and ARPE-HPV eGFP-CLCa+GAK(AP2*)-mRuby cells (#4 from left) that were treated with GAK siRNA. Images were obtained at 1 frame/s and collected for 7.5 min. Movie is accelerated 25-fold.

Movie 3: Time-lapse TIRFM imaging of ARPE-HPV eGFP-CLCa+GAK(WT)-mRuby cells (left), ARPE-HPV eGFP-CLCa+GAK(clathrin*)-mRuby cells (middle) and ARPE-HPV eGFP-CLCa+GAK(J*)-mRuby cells (right) that were treated with GAK siRNA. Images were obtained at 1 frame/s and collected for 7.5 min. Movie is accelerated 25-fold.

### Supplemental Figures

**Figure S1.**
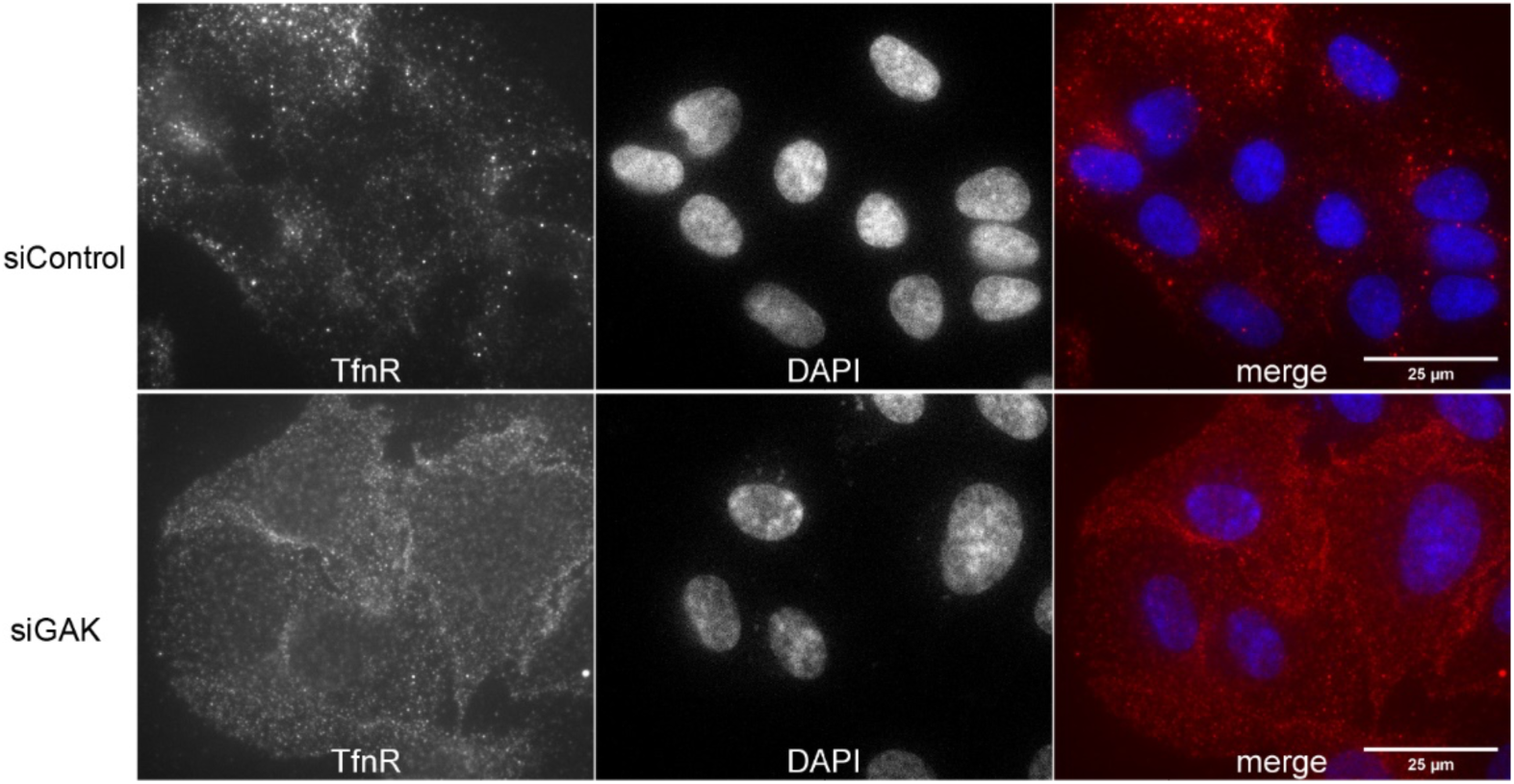
GAK knockdown leads to the accumulation of TfnR on the surface of ARPE-HPV eGFP-CLCa cells. Images shown were acquired by immunofluorescence and TIRFM imaging. Scale bars = 25 µm.

**Figure S2.**
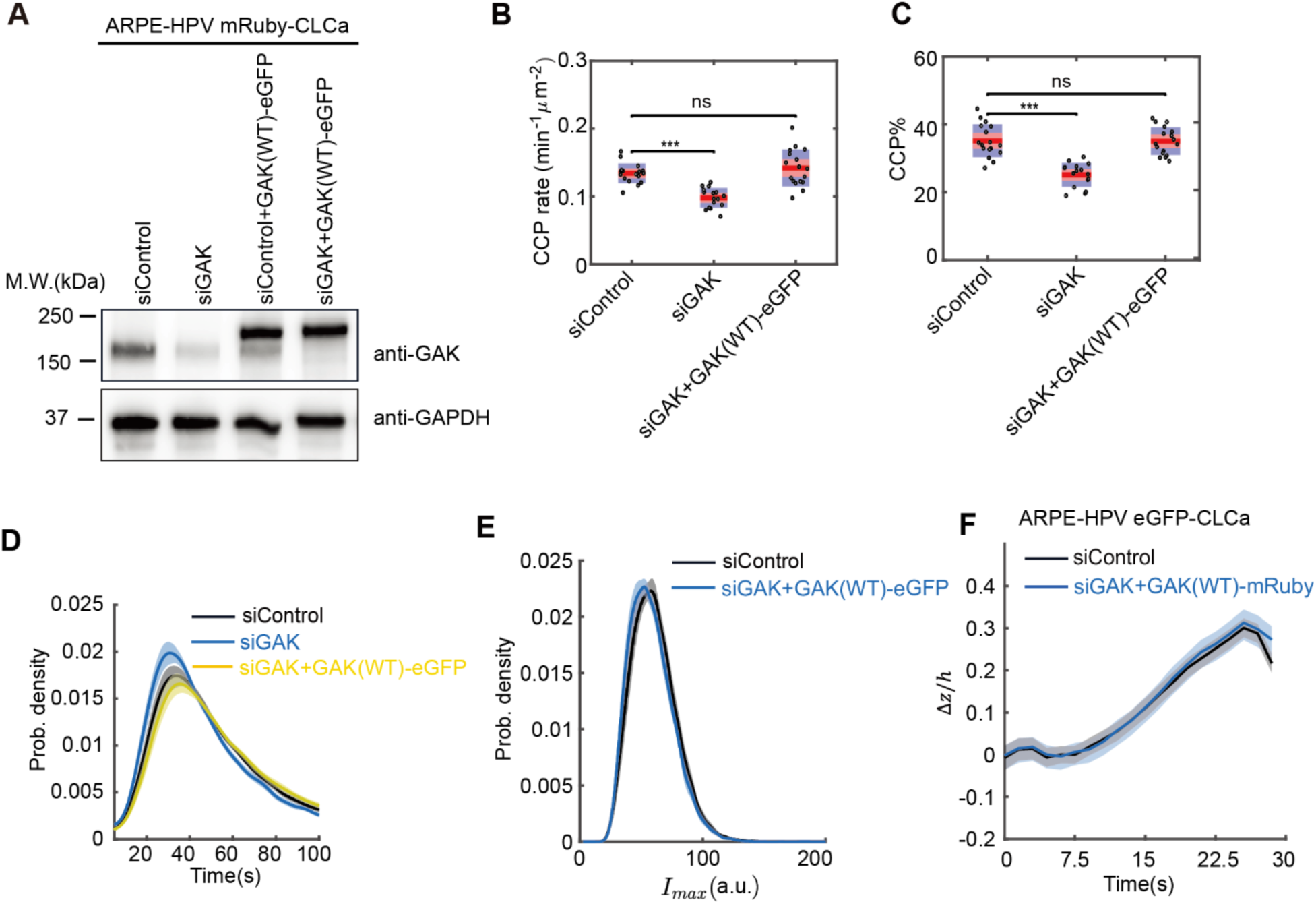
Expression of GAK-eGFP(WT) or GAK-mRuby(WT) rescues the phenotypes of GAK knockdown. **(A)** Western blotting indicates the expression level of exogenous GAK-eGFP in ARPE-HPV mRuby-CLCa cells and the efficiency of 3’ UTR siRNA-mediated knockdown of GAK. **(B-C)** Expression of GAK(WT)-eGFP rescued the reduction in the initiation rate and % of CCPs induced by GAK knockdown. Statistical analysis of the data in (B) and (C) is the Wilcoxon rank sum test, ns: P > 0.05; ***: P<0.001. **(D-F)** Expression of GAK(WT)-eGFP restored (D) the lifetime distribution, (E) the maximum fluorescence intensity distribution, and (F) the invagination depth of bona fide CCPs to control values. Data presented were obtained from n = 15 movies for each condition. Each dot in (B,C) represents a movie. Number of dynamic tracks analyzed IN (B-E): 180838 for siControl, 185121 for siGAK, 187323 for siGAK+GAK(WT)-eGFP. Number of CCP tracks analyzed to obtain the Δ*z*/*h* curves in (F): 15,395 for siControl, 10,880 for siGAK+GAK(WT)-mRuby. Shadowed area in (D-F) indicates 95% confidential interval.

**Figure S3.**
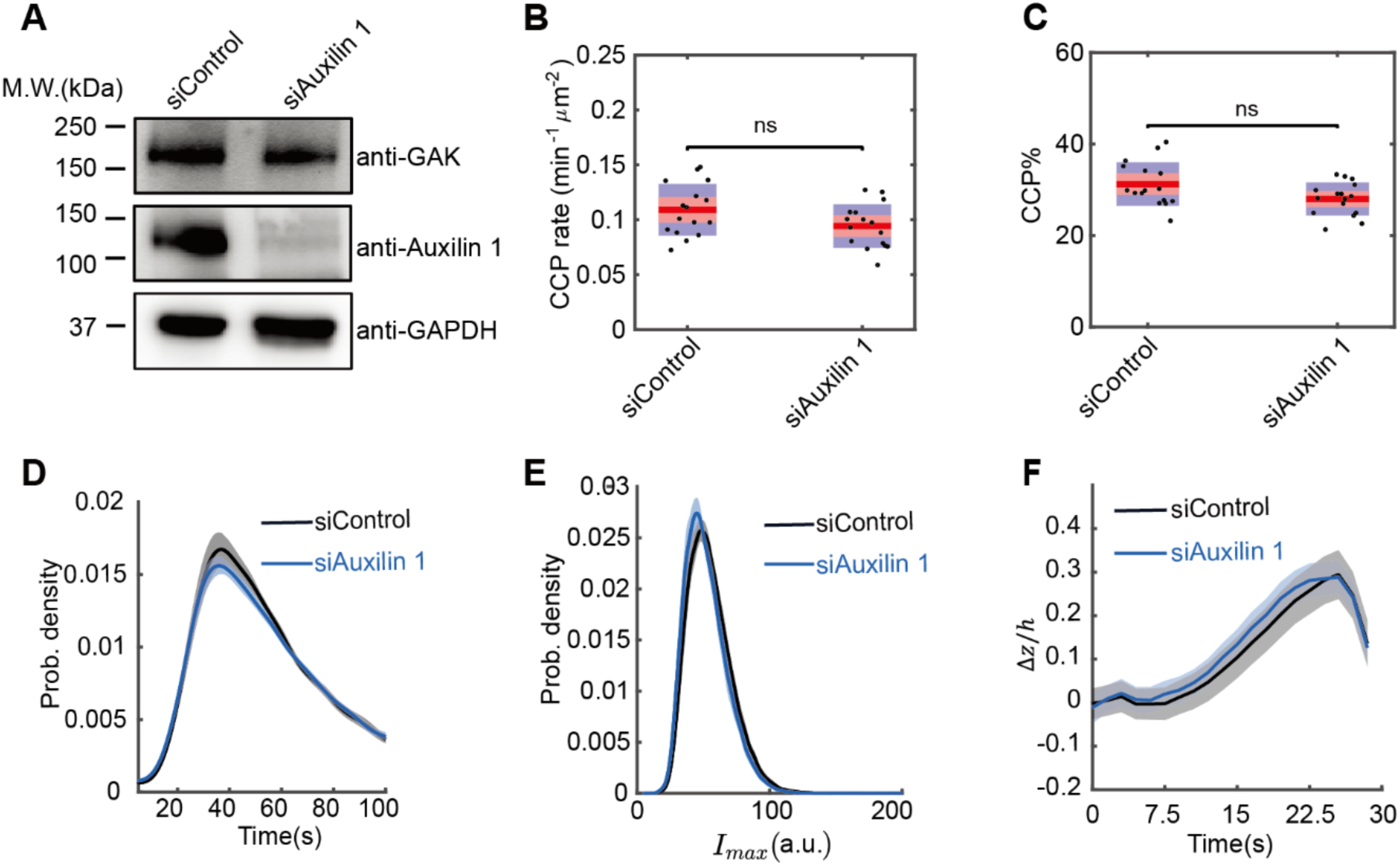
Auxilin Knockdown does not affect CCP formation or invagination. **(A)** siRNA-mediated knockdown efficiency of Auxilin in ARPE-HPV eGFP-CLCa cells probed by western blotting. Auxilin knockdown did not affect GAK levels. **(B-C)** Auxilin 1 knockdown did not affect the initiation rate and % of bona fide CCPs. Statistical analysis of the data in (B) and (C) is the Wilcoxon rank sum test, ns: P > 0.05. **(D-F)** Auxilin knockdown did not affect (D) the lifetime distribution, (E) the maximum fluorescence intensity distribution, or (F) the invagination depth of bona fide CCPs. Data presented were obtained from n = 15 movies for each condition. Each dot represents a movie. Number of dynamic tracks analyzed:166,745 for siControl, 149,623 for siAuxilin 1. Number of CCP tracks analyzed to obtain the Δ*z*/*h* curves in (F): 16,972 for siControl, 8,332 for siAuxilin 1. Shadowed area in (D-F) indicates 95% confidential interval.

**Figure S4.**
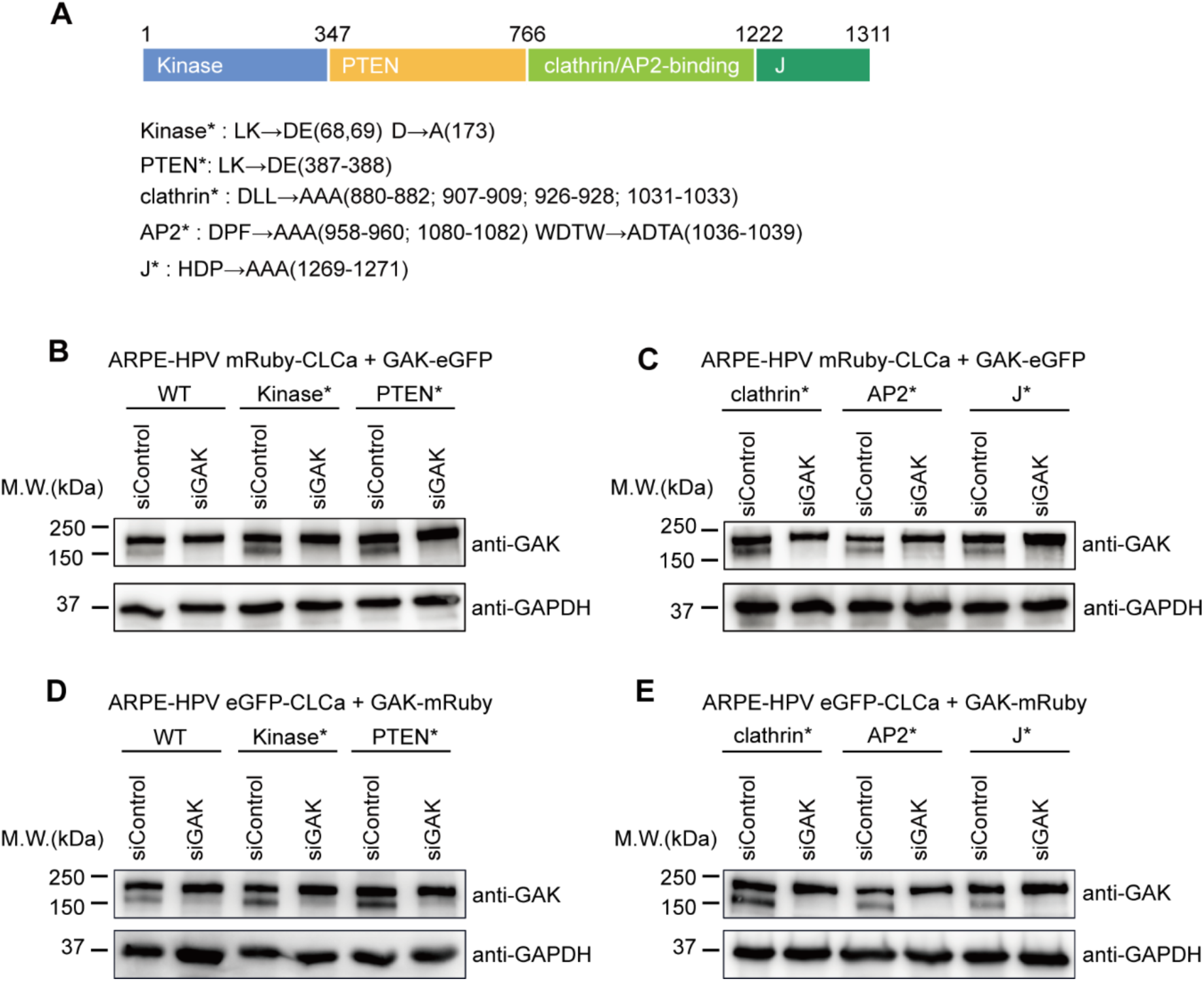
Generation of GAK mutants and stable cell lines. **(A)** Domain structure of GAK and corresponding mutations (denoted with *) to abolish domain functions. **(B-C)** Western blotting of ARPE-HPV mRuby-CLCa cells that stably express GAK-eGFP (WT and mutant constructs) and treated with control siRNA or GAK siRNA. Treatment of GAK siRNA knocked down endogenous but not exogenous GAK. **(D-E)** Western blotting of ARPE-HPV eGFP-CLCa cells that stably express GAK-mRuby (WT and mutant constructs) and treated with control siRNA or GAK siRNA. Treatment of GAK siRNA knocked down endogenous but not exogenous GAK.

**Figure S5.**
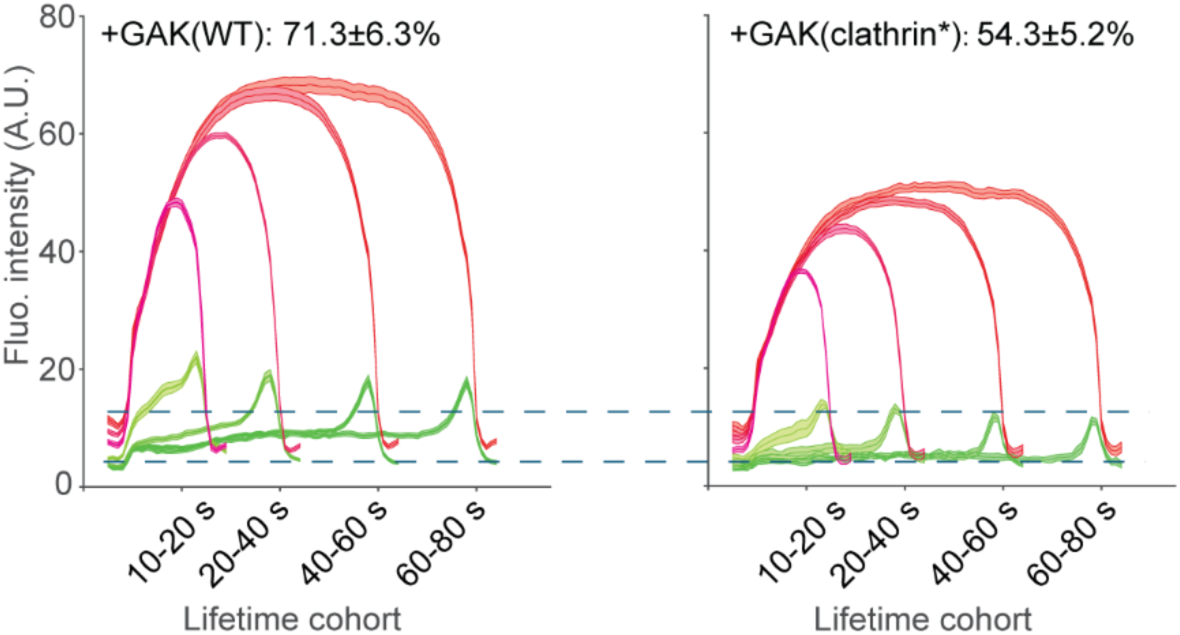
Recruitment of GAK(clathrin*) to CCPs is inhibited but not abolished when compared with GAK(WT). Fluorescence intensity cohorts of mRuby-CLCa (red) and GAK-eGFP (green) were generated using dual-channel TIRFM imaging and cmeAnalysis. GAK(WT) was recruited to 71.3±6.3% CCPs while GAK(clathrin*) was recruited to 54.3±5.2% CCPs. In addition,the magnitude of GAK(WT) recruitment was higher than that of GAK(clathrin*).

**Figure S6.**
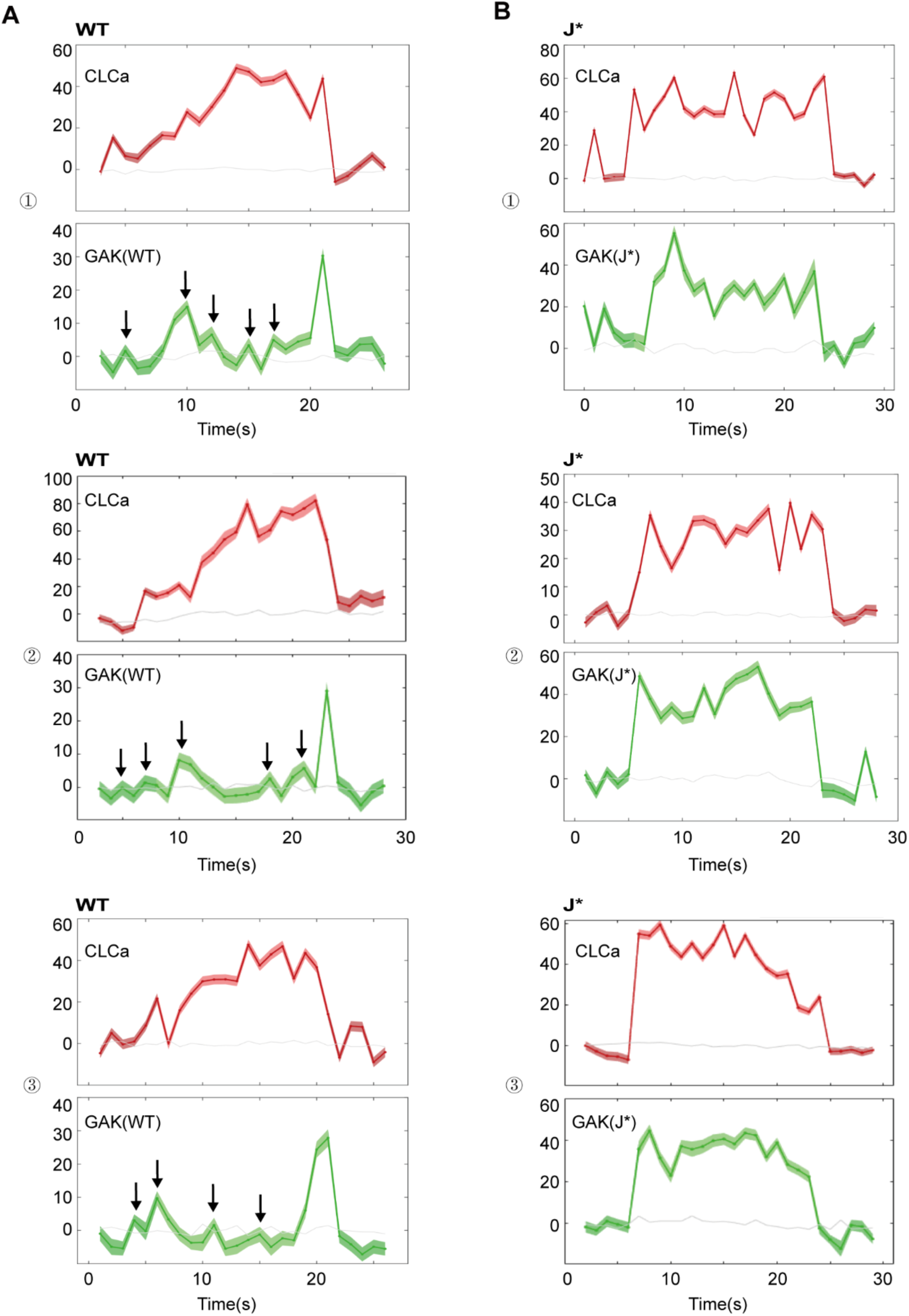
Additional representative dual-channel (mRuby-CLCa and eGFP-GAK) tracks of CCPs from ARPE-HPV mRuby-CLCa cells that stably express GAK-eGFP: **(A)** WT and **(B)** J*.

**Table S1:**
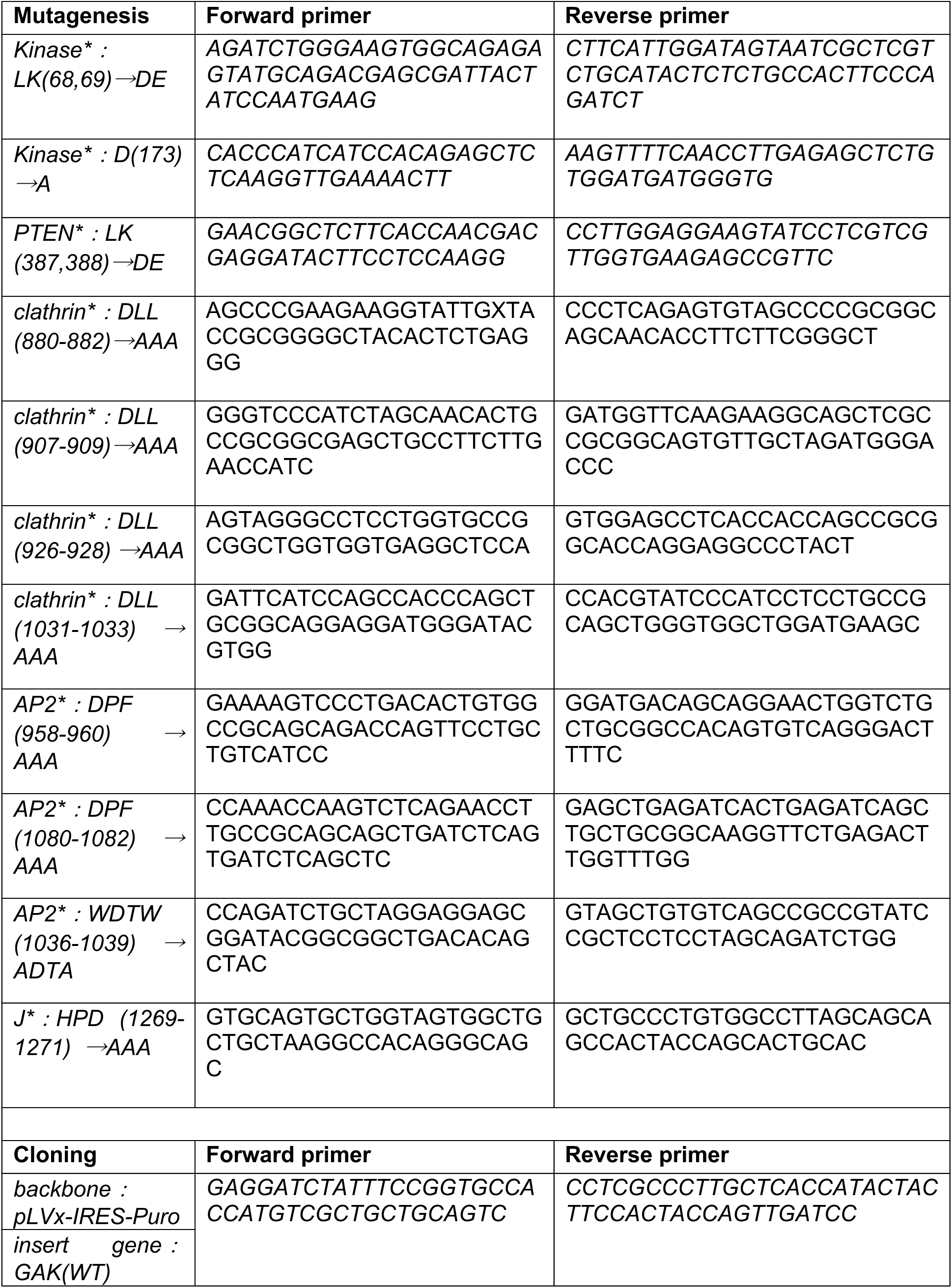
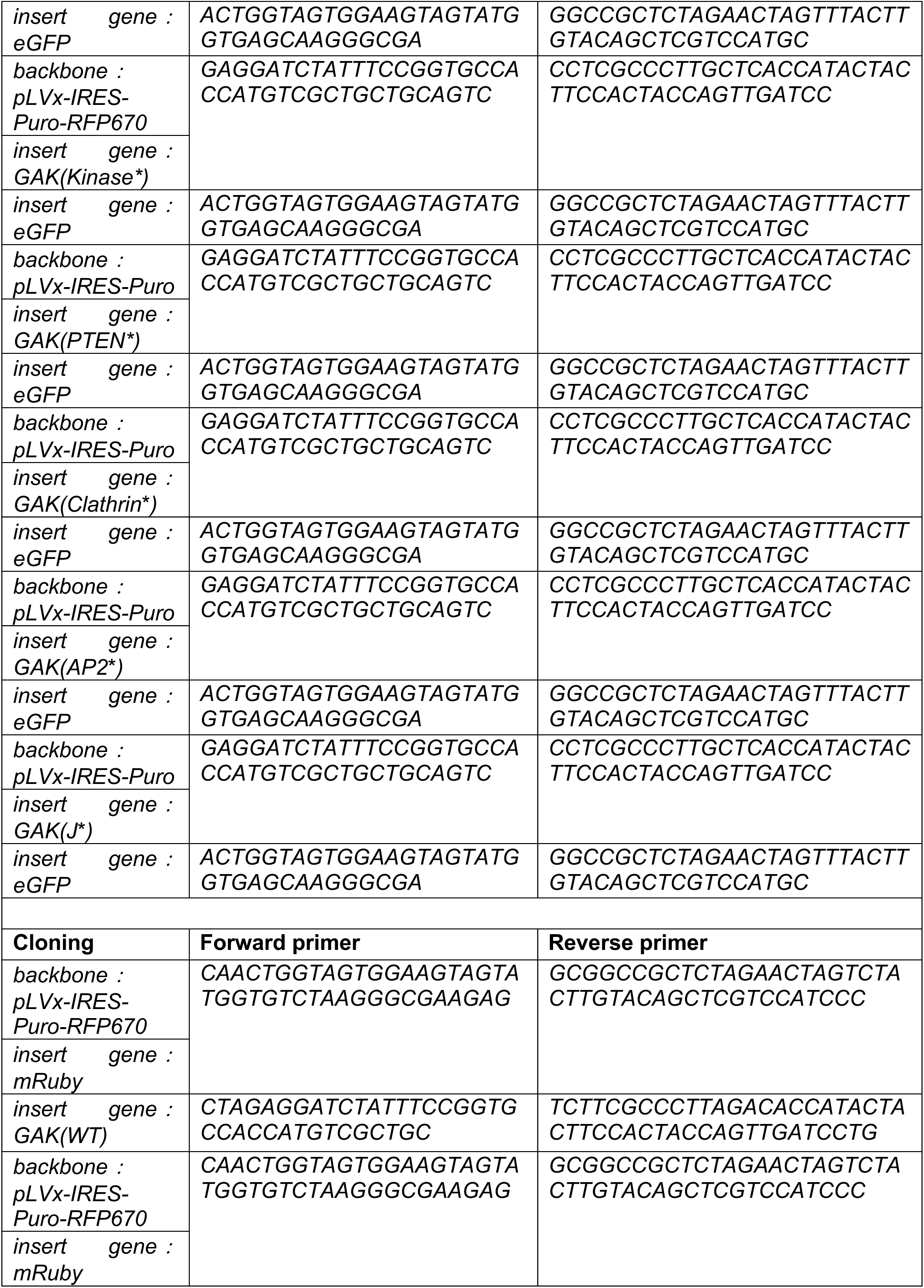
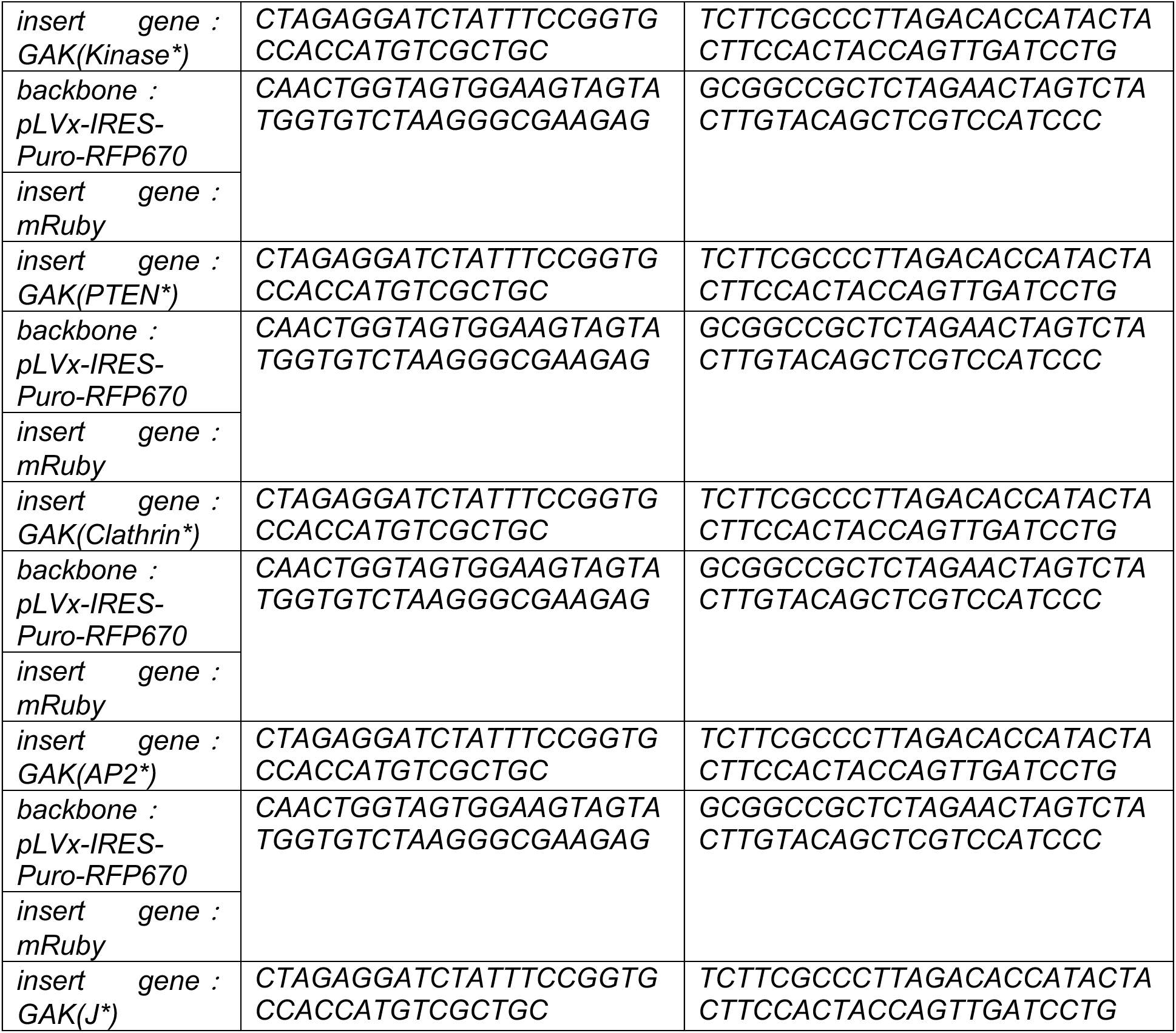
List of primers.

## Notes

### Competing Interest Statement

The authors have declared no competing interest.

